# Rapid Population Decline of the Pillar Coral *Dendrogyra cylindrus* Along the Florida Reef Tract

**DOI:** 10.1101/2020.05.09.085886

**Authors:** Karen L. Neely, Cynthia L. Lewis

## Abstract

Coral reefs worldwide are in a state of decline, but the status of populations and stressors for rare species are generally not well documented using broad-scale monitoring protocol. We fate-tracked all known colonies of the pillar coral *Dendrogyra cylindrus* on the Florida Reef Tract from 2013 – 2020 to assess the population and document the impacts of chronic and acute stressors. Large average colony size, an absence of juveniles, and large geographic distances between genotypes suggest that the Florida *D. cylindrus* population has been reproductively extinct for decades. During the study period, low-intensity chronic stressors were balanced by regrowth, while back-to-back years of coral bleaching and thermally-exacerbated disease led to declines that the subsequent years of recovery suggest would take 11 uninterrupted years to overcome. The most recent stressor on Florida’s *D. cylindrus* population is “stony coral tissue loss disease.” Following the appearance of the disease in Florida in 2015, it resulted in unrecoverable losses to the *D. cylindrus* population as tissue, colonies, and whole genotypes were driven to extinction. Losses of 91% of coral tissue, 88% of colonies, and 73% of genotypes between 2014 and early 2020 have led to functional extinction of *D. cylindrus* on the Florida Reef Tract.

## Introduction

The pillar coral *Dendrogyra cylindrus* is a conspicuous stony coral described as uncommon or rare in reef environments throughout the Caribbean (Szmant 1986). It is taxonomically unique as the only member of its genus, structurally unique as the only Caribbean columnar coral, and behaviorally unique as a species whose polyps are extended during light as well as dark hours.

*Dendrogyra cylindrus* is listed as Vulnerable on the IUCN Red List (Aronson et al. 2008) and Threatened under the US Endangered Species Act (USFWS 2014). It was also listed as Threatened by the state of Florida in 2011 and, like all stony corals on the Florida Reef Tract, is protected under Florida’s Coral Reef Protection Act and the Florida Fish and Wildlife Conservation Commission’s Marine Life Rule, which prohibits take, injury, or destruction of stony corals. However, population information on *D. cylindrus* throughout its range has been scarce, and with one exception in Colombia (Bernal-Sotelo et al. 2019), population assessments, stressors, and trends have remained largely undocumented.

The Florida Reef Tract (FRT) represents the northernmost extent of most Caribbean reef corals. Declines in coral cover over decadal scales on this Florida barrier reef system are well documented (Porter and Meier 1992; Ruzicka et al. 2013). Many of these losses are the result of short-term stressors such as bleaching or disease events causing the demise of long-lived colonies, often decades to centuries old. Decades of acute stressors on the Florida Reef Tract include historic bleaching events (Causey 2008), outbreaks of white band disease in Acroporids (Aronson and Precht 2001), and outbreaks of white plague, black band, and yellow-band diseases on a wide range of species (Goreau et al. 1998; Richardson et al. 1998; Sutherland et al. 2004). Chronic declines in health related to predation (Baums et al. 2003) and water quality (Thurber et al. 2014; Lapointe et al. 2019) have also occurred. Beginning in 2014, a series of high-intensity stressors included two consecutive hyperthermal bleaching events (summer 2014 and summer 2015) and the outbreak of a temporally and geographically extensive disease termed “stony coral tissue loss disease” (SCTLD) (Precht et al. 2016; FKNMS/DEP 2018).

The impact of stressors may vary by event, region, and species, but declines in rare species are more likely to lead to reproductive isolation or extinction (Courchamp et al. 1999; Chan et al. 2019). Population status of rare species is also more difficult to assess because traditional survey techniques are ineffective at capturing them. *Dendrogyra cylindrus,* for example, is seldom seen in randomized survey techniques in Florida and must be systematically targeted to attain quantitative information. Such targeting allows for fate-tracking of individuals in order to document status and trends over time and through various environmental events.

This study assessed the population demographics of *D. cylindrus* colonies on the FRT and documented their fate through chronic and acute stressors from 2014 – 2020. It followed the decline of a rare species into near-extinction in the region.

## Methods

### Groundtruthing and Monitoring

Beginning in 2013 – 2014, potential locations of *D. cylindrus* colonies on the FRT were collated using databases from various monitoring and outplanting projects as well as local knowledge from dive shops and other resource users. From October 2013 to May 2014, an area approximately 700 m^2^ surrounding each provided coordinate was searched for the presence of *D. cylindrus* colonies. A total of 543 colonies were identified during this time period. From June 2014 through March 2020, an additional 272 colonies were identified and surveyed. Most data for the 68 colonies north of Biscayne National Park in the Southeast Florida region are derived from work conducted by the Gilliam lab (Nova Southeastern University) and published in Kabay (2016). At each site, corals were identified with cattle tags or by maps and photographs so that each individual could be accurately identified in future surveys.

In spring of 2014, a subset of 42 geographically stratified sites were selected for tri-annual monitoring to document trends and seasonal changes. These sites included 27 individual isolated colonies, five sites with 2 – 3 colonies, and ten sites with 4 or more colonies. At three sites with greater than 50 colonies, a subset of 24 colonies were selected for fate-tracking. These 24 colonies were selected to spatially cover the site and to incorporate both large and small colonies. At all other sites, all colonies were monitored. Sites were selected to span the geographical range of the species along the FRT and to represent a variety of depths. At ten of these sites, data loggers (Onset HOBO Inc., Bourne, Massachusetts USA) placed at the base of a colony recorded hourly temperatures. Additional data on thermal stress were acquired using NOAA’s Coral Reef Watch indices of degree heating weeks, which sum thermal stress (number of degrees Celsius above the highest summertime mean sea surface temperature) across rolling 12-week periods (NOAA 2013-2019).

The subset of 42 sites were visited tri-annually in spring (April-May), fall (September-October), and winter (January-February) from spring 2014 to spring 2017 for a total of nine sampling periods. All known colonies were re-visited in spring of 2017 and also in spring of 2018 for full colony assessments. Other monitoring occurred opportunistically from 2017-2020.

### Genetic Diversity

Full sampling and analysis protocols for determining genetic variability within FRT *D. cylindrus* is presented in Chan et al. (2019). Briefly, tissue clips and/or syringe tissue biopsy samples (Kemp et al. 2008) were taken from 161 colonies across within the Florida Reef Tract. Sample selection was stratified by depth and by the number of colonies at the sites. Microsatellites were designed to identify host and symbiont diversity. Results indicated instances of genetic clonality as well as uniqueness within and across sites. The term “colony” is used to define any individual distinctly separated from other colonies by no apparent skeletal connectivity; the term “ramet” is used to define colonies that were asexually reproduced and are genetically identical to other surrounding ramets.

Utilizing these results, the number of genotypes across the Florida Reef Tract was conservatively estimated by clustering non-genotyped colonies within the minimum distance (70 m) identified as clonal by Chan et al. into a single genotype. Non-genotyped outside of that distance from a nearest neighbor were classified as unique genotypes.

### Demographics

Linear measurements of maximum length, width, and height of each colony were taken with a 50 cm measuring stick. Surface area was estimated as that of a half ovoid using length, width, and height measurements as axes; live surface area was then calculated by multiplying that by the percent live tissue on the colony. Tissue gains or losses over time were calculated by subtracting the live surface area at each monitoring event from the live surface area at the baseline. Volume estimates were calculated as a half ovoid using the same axes measurements. Depth at the base of each colony was identified using a dive computer.

For population analyses on nearest neighbors, depth, and size, only the largest colony from each known or presumed genotype was used. It is presumed that the largest colony best represents the original recruited coral, with all smaller ramets resulting from stochastic fragmentation events.

### Health Status and Mortality

During each monitoring event, the percent of each colony covered with live tissue, old mortality, and recent mortality (classified as bare white skeleton with defined polyp structure) was recorded in the field. The vast majority of percentage assessments were done by the same diver, but in a few instances an alternate diver recorded percentages. Comparative assessments between the two divers were done periodically and found to be in agreement. When recent mortality was observed, the cause(s) were determined and recorded to the extent possible. If multiple stressors had caused mortality, the percent attributable to each was differentiated. Bleaching/paling was also recorded categorically as “paling,” “partial bleaching,” and “bleaching.”

The number of observations of each stressor was compared to the total number of *D. cylindrus* monitoring records to identify prevalence. The amount of recent mortality attributed to each stressor was averaged to quantify the severity.

### Analyses

Using GIS layering, colony locations were overlaid on the Disturbance Response Monitoring zone map (FRRP 2015) to analyze inshore to offshore reef distribution. Colonies were also classified to habitat by overlaying locations on the Florida Unified Reef Map (FWRI 2014).

Differences in size metrics between regions of Florida and between the Florida population and other Caribbean studies were assessed using t-tests or ANOVAs when normality and equal variance were met, and by using Kruskal-Wallis ANOVA on Ranks when they were not. Size metrics and depth were compared using a linear regression. Comparisons of stressor prevalence between regions were conducted using z-tests. All analyses of significance are reported with α = 0.05.

## Results

### Population Distribution

Between 2013 and March 2020, a total of 815 *Dendrogyra cylindrus* colonies were recorded along the Florida Reef Tract (Figure 1). Of these total known colonies, 11 were within Dry Tortugas National Park, 709 were within the Florida Keys National Marine Sanctuary (FKNMS), 27 were within Biscayne National Park, and 68 were north of Biscayne National Park in the Southeast Florida region. Management zones within these regions offered varying levels of resource protection and usage patterns. Of the 815 colonies, 144 (18%) were within no-take marine protected areas (143 in FKNMS Sanctuary Preservation Areas, and 1 within the Dry Tortugas Research Natural Area), 36 (4%) were within zones that limit fishing to catch-and-release trolling, and 635 (78%) were in fishable waters.

**Figure 1.**
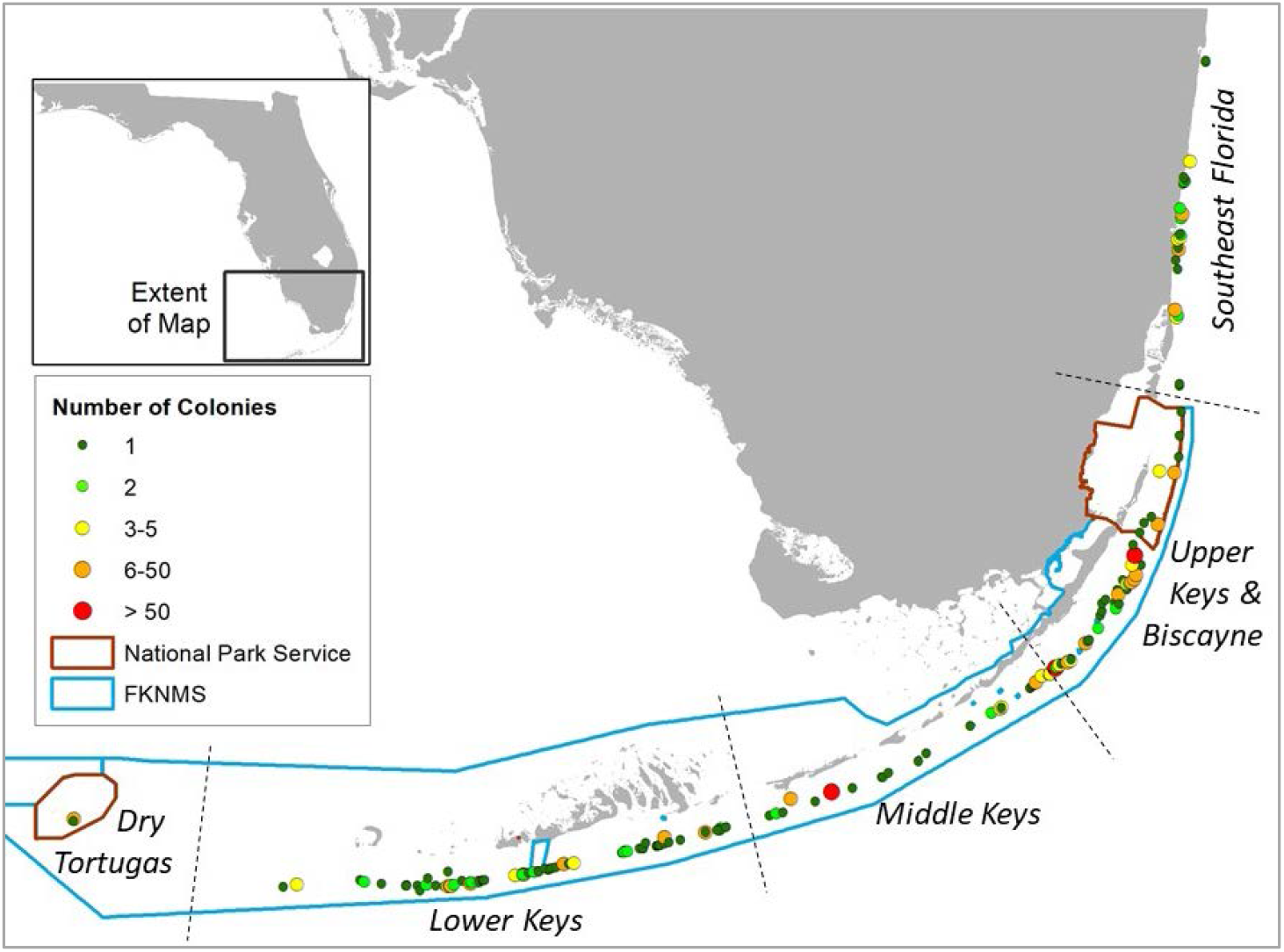
Location of *Dendrogyra cylindrus* colonies along the Florida Reef Tract. Color/size differences represent the number of colonies at each site. National Park and Florida Keys National Marine Sanctuary boundaries are shown.

Colony depth averaged 7.4 (±2.5 SD) meters. The shallowest site was 2.4 meters and the deepest was 19.5 meters (Figure 2). The two deepest colonies were at the two northernmost sites.

**Figure 2.**
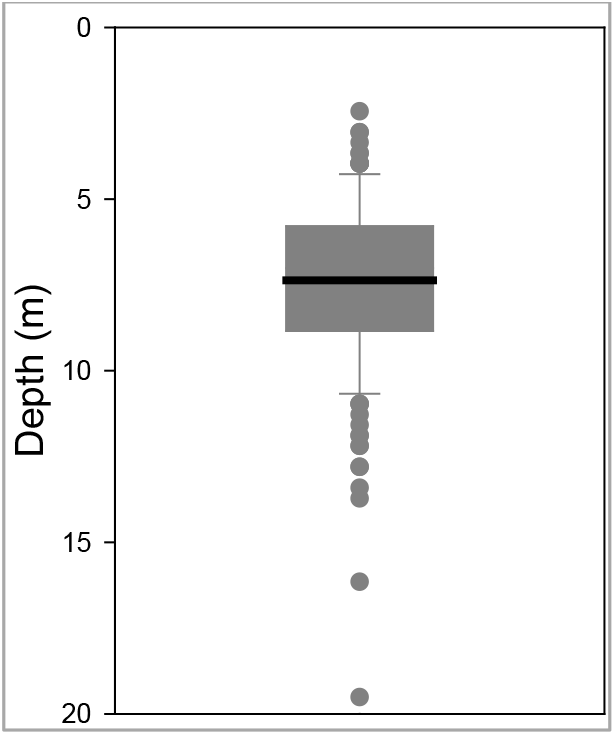
Box plot of *D. cylindrus* colony depth on the Florida Reef Tract. For genotypes with multiple ramets, the depth of only the largest ramet was included. Box shows the quartiles with the mean. Whiskers incorporate the 10^th^ – 90^th^ percentiles. Dots are outliers.

Within the Southeast Florida region, the DRM map delineated inshore, inner reef, middle reef, and outer reef zones. *D. cylindrus* colony distribution on these was: 21 inshore, 40 inner reef, 3 middle reef, and 2 outer reef (Figure 3a). Within the Biscayne to Dry Tortugas area, the map classified mid-channel patch reef, offshore patch reef, and forereef areas. *D. cylindrus* colony distribution on these was 20 mid-channel patch reef, 82 offshore patch reef, and 648 forereef (Figure 3b). Overlays from the Unified Reef Map identified 339 colonies located on spur and groove reef, 111 on patch reef/isolated reef structures, 13 on ridges, and 314 on low-relief contiguous reef/colonized pavement (Figure 3c).

**Figure 3.**
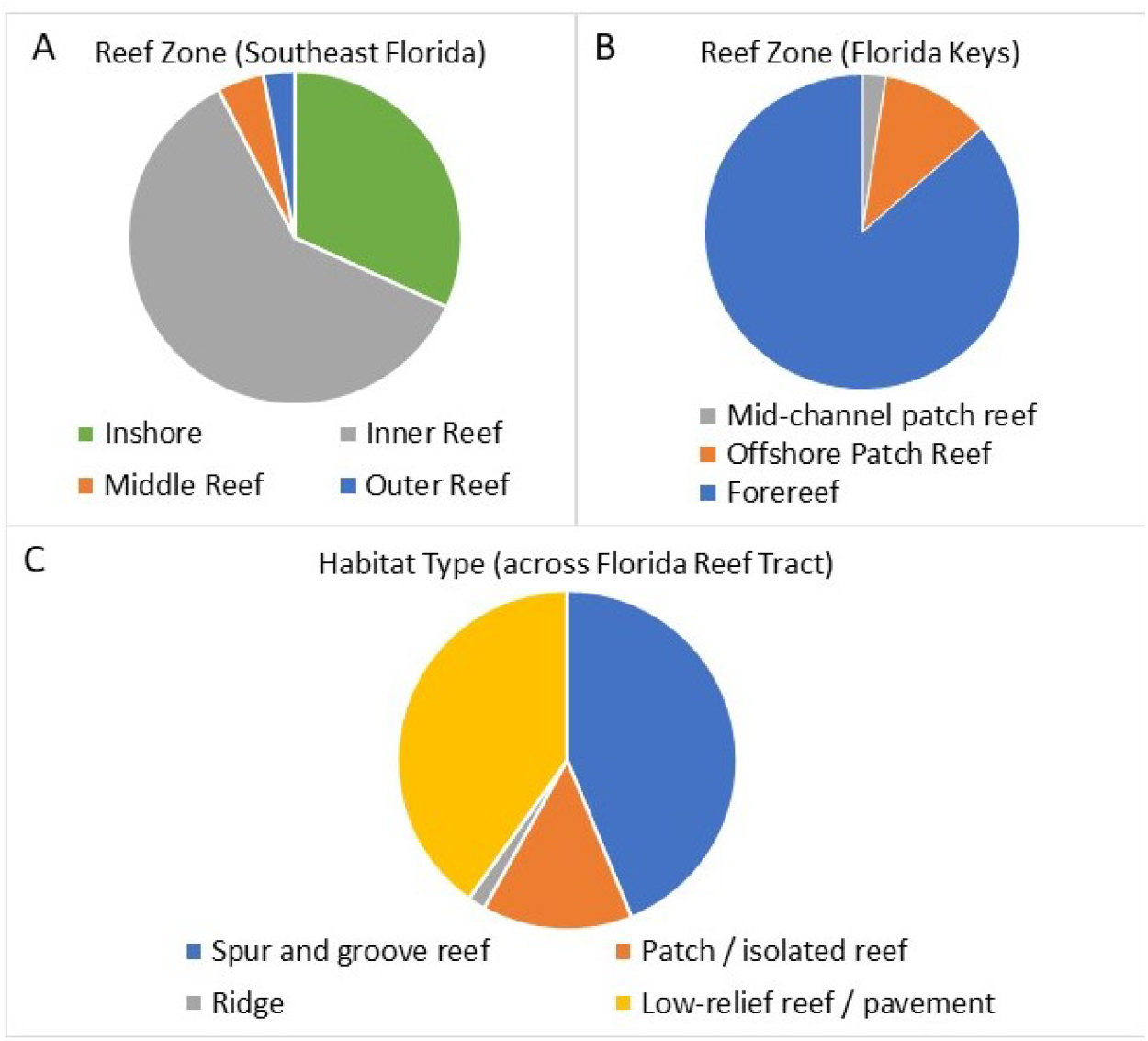
*Dendrogyra cylindrus* zone and habitat distribution. Maps from the Disturbance Monitoring Program identified reef zone distribution of colonies within southeast Florida (A) and the Florida Keys (B). The Florida Unified Reef Map identified the nearest habitat type across the whole of the Florida Reef Tract (C).

### Genetic Diversity

Genetically sampled *D. cylindrus* colonies identified clonality at most sites containing multiple colonies (Chan et al. 2019). The maximum distance between ramets in Florida was identified at Pickles Reef (Upper Keys) where 110 colonies were distributed across a maximum diameter of 84 meters. Three other sites had distances between individual ramets that exceeded 60 meters; the greatest was 69.9 meters.

In order to estimate the number of genotypes within the Florida Reef Tract to include colonies that were not genetically sampled, all unsampled colonies or groups of colonies that were within 70 meters of other colonies were presumed to be of the same genotype. All colonies or groups of colonies that were greater than 70 meters from any other colonies were presumed genetically distinct. These assumptions resulted in an estimated 188 unique genotypes on the Florida Reef Tract.

Colony numbers were not evenly distributed across the genotypes. More than 50% of Florida colonies were represented by only 5 genotypes (Figure 4). Of the 188 genotypes, 62% (117) were represented by a single colony each. Overall, asexual reproduction accounted for 77% (627/815) of Florida *D. cylindrus* colonies.

**Figure 4:**
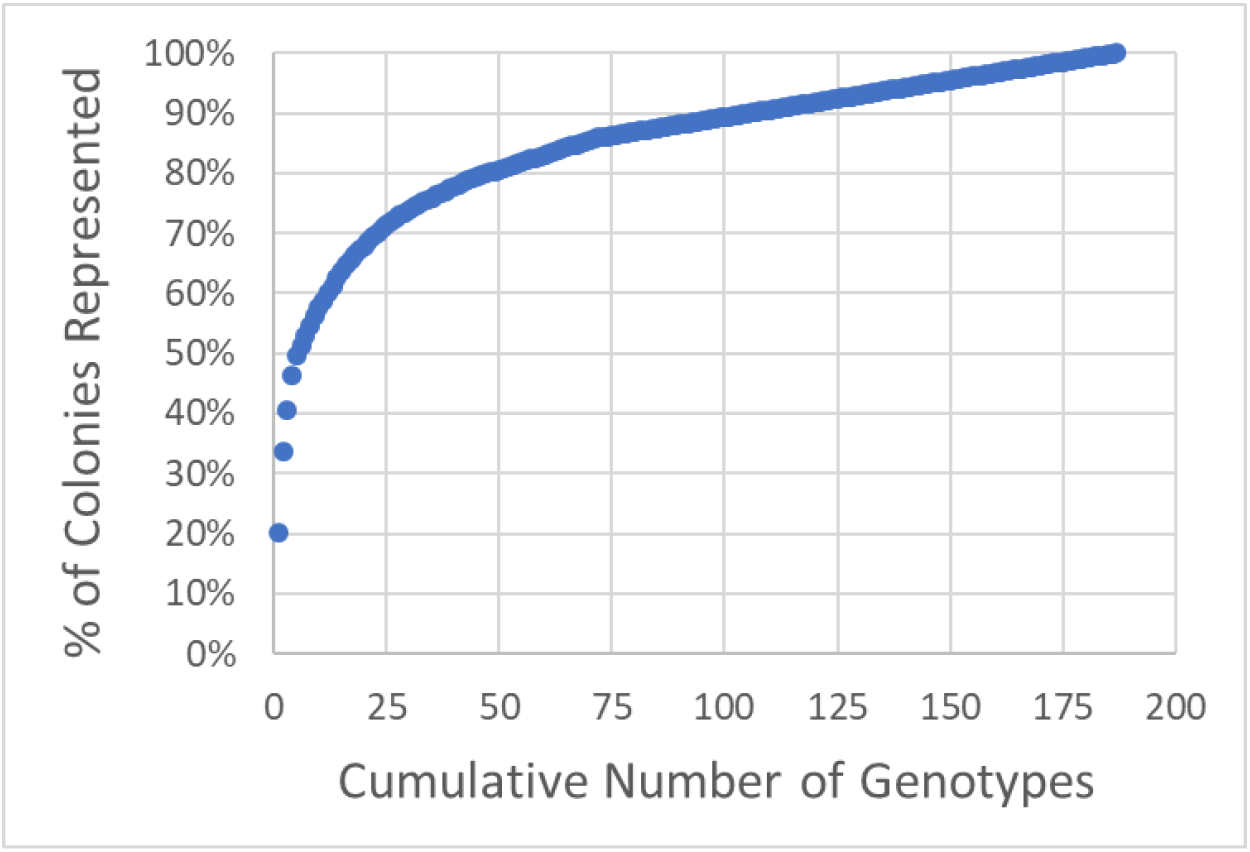
Cumulative curve showing the number of Florida *Dendrogyra cylindrus* colonies represented by distinct genotypes. Over 50% of colonies were accounted for by the five most populous genotypes.

Distance between genotypes ranged from 2.5 to 6596 meters. The average distance was 1070 (± 1367 SD) meters (Figure 5). Of the 188 distinct genotypes, 37% (70) were greater than 1 km from their nearest neighbor, 40% (76) were between 100-1000 m, 20% (38) were between 10 and 100 m, and 2% (4) were less than 10 m.

**Figure 5.**
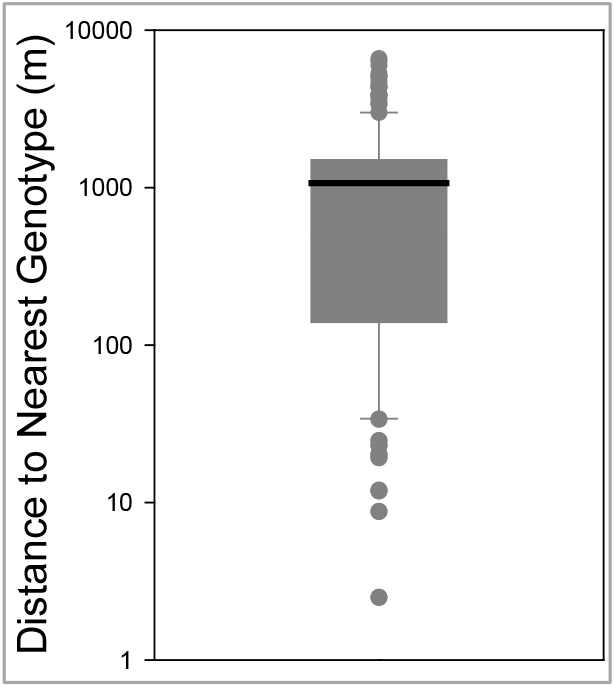
Box plot of logarithmic distances between *Dendrogyra cylindrus* genotypes. Box shows the quartiles with the mean (1070 m). Whiskers incorporate the 10^th^ – 90^th^ percentiles. Dots are outliers.

### Demographics

The size of the largest *D. cylindrus* colony within each genotype was used to assess demographics. Colony size averaged 167 cm in diameter and 123 cm in height. Estimated surface area averaged 15.6 m^2^ and estimated volume averaged 10.9 m^3^ (Table 1). The largest colonies exceeded 5 meters in length, 3 meters in height, and 114 m^3^ in volume. In 2014, a single juvenile colony with a diameter of 7 cm was found in the Upper Keys, but had perished within a year of first siting.

**Table 1.** Size metrics for *Dendrogyra cylindrus* colonies on the Florida Reef Tract. Minima, averages (± standard deviations), and maxima are shown for maximum diameter, height, surface area, and volume. Surface area and volume are derived from formulae for half a spheroid using length, width, and height dimensions. For genotypes with multiple colonies, only the largest colony was used in analyses.

|  | Diameter (cm) | Height (cm) | Surface area (m<sup>2</sup>) | Volume (m<sup>3</sup>) |
| --- | --- | --- | --- | --- |
| Minimum | 7 | 1 | 0.01 | 0.0002 |
| Average ± SD | 168 ± 89 | 123 ± 66 | 15.6 ± 15.3 | 10.9 ± 16.0 |
| Maximum | 531 | 300 | 89.3 | 114.9 |

No measure of size (maximum diameter, height, surface area, volume) correlated significantly with depth (linear regressions: p > 0.13). However, size differences were observed within different regions of the Florida Reef Tract. The average volumetric size of *D. cylindrus* colonies in the Lower Keys was 8.0 m^3^, while average size in all other regions exceeded 10 m^3^ (Figure 6). Volumetric distributions did not pass Shapiro-Wilk normality, and ANOVA on Ranks determined differences between regions were not significant (p = 0.2). However, when Lower Keys colonies and Upper Keys colonies were compared, volumetric differences were significant (Kruskal-Wallis ANOVA on Ranks: p = 0.04.

**Figure 6.**
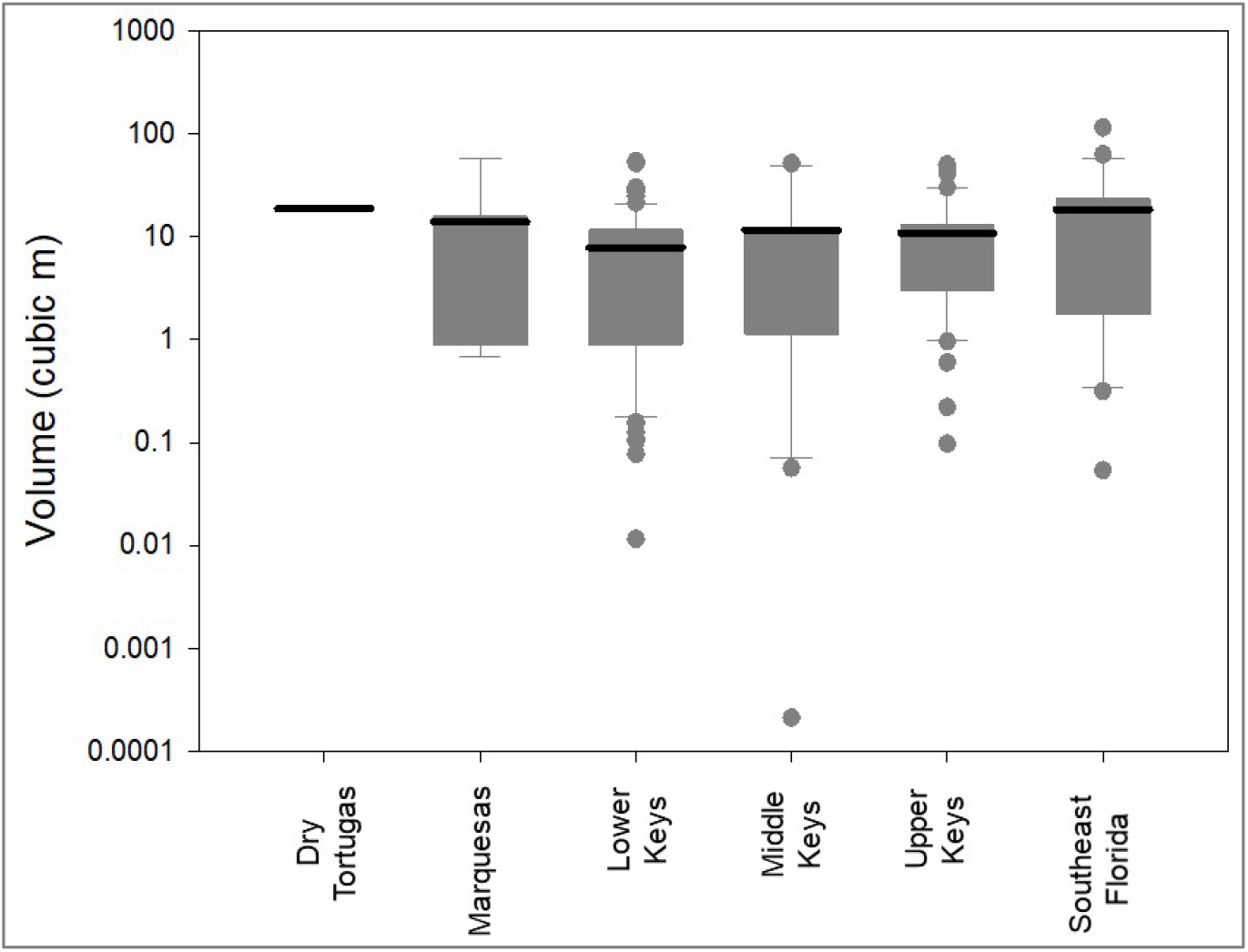
Box plots (logarithmic) of *Dendrogyra cylindrus* colony volumetric estimates separated by reef region. Boxes show the quartiles with the mean. Whiskers incorporate the 10^th^ – 90^th^ percentiles. Dots are outliers. Size analyses include only the largest colony within each genotype.

### Baseline Status

During baseline surveys in 2013-2014, a total of 542 colonies were identified and surveyed; 533 of these were alive. The average live coral cover on all colonies (including the dead ones) was 70%, and the average recent mortality was 0.5%. Twenty two percent (117/542) of the baseline colonies exhibited recent mortality. On those, the average recent mortality was 2.2%.

From 2015 to March 2020, an additional 273 colonies were identified and surveyed, increasing the total known colonies to 815. If we parsimoniously assume no gains or losses in live tissue from the time of “new” colony identification back to the baseline surveys, the total amount of live *D. cylindrus* tissue estimated across the Florida Reef Tract in 2013 – 2014 was 2767 m^2^. However, this estimate is conservative, as most colonies identified from 2015 onwards had already suffered mild to extensive mortality.

### Minor and Chronic Sources of Mortality

*Dendrogyra cylindrus* colonies were subject to a variety of minor, chronic sources of mortality including damselfish gardens and nests, predation by the corallivorous snail *Coralliophila abbreviata*, competition with other benthic organisms, and physical damage from abrasion or burial.

Out of 3065 observations on live colonies, 28 observations (< 1 %) recorded recent mortality from damselfish gardens (primarily from the threespot damselfish *Stegastes planifrons*) or damselfish nests (primarily from the sergeant major *Abudefduf saxatilis*). Recent mortality on colonies affected by these gardens or nests averaged 0.6%. Mortality as a result of damsel gardens was observed year-round, but mortality associated with damsel nests was seen only in March through May (Figure 7).

**Figure 7.**
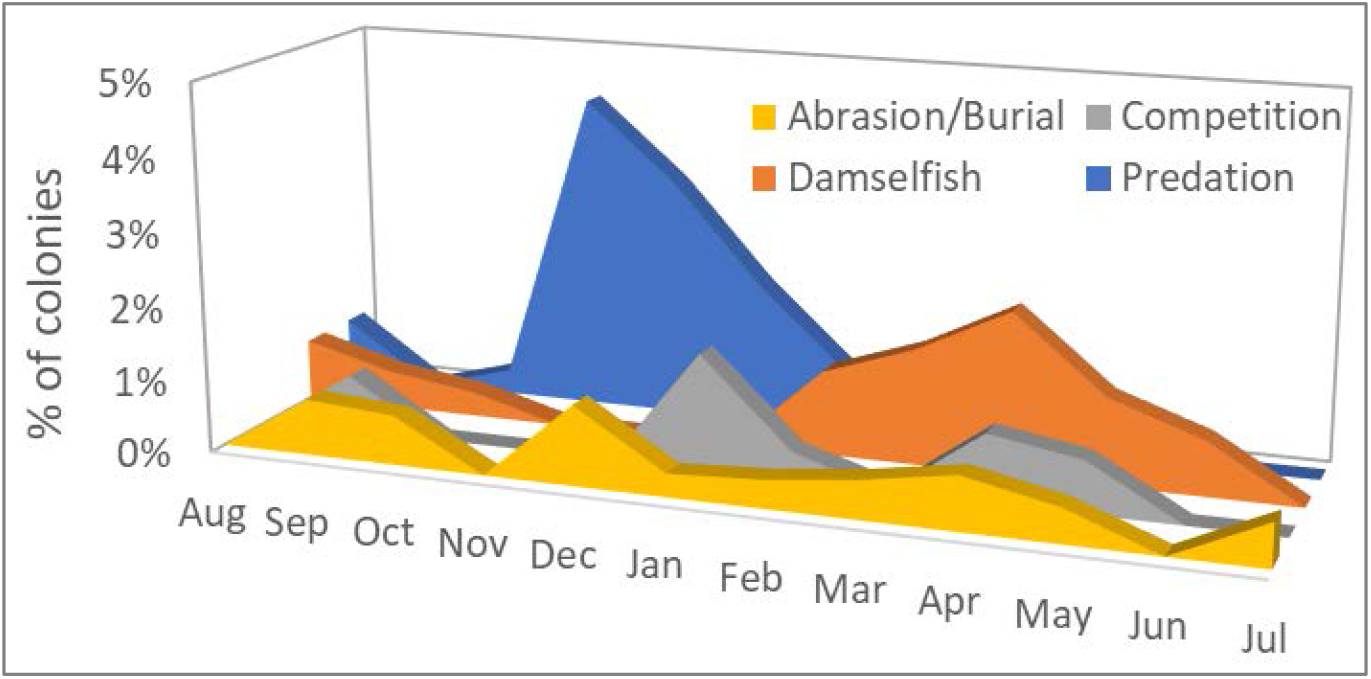
Percentage of observations on live *Dendrogyra cylindrus* colonies in which recent mortality could be attributed to four chronic stressors: abrasion/burial, competition with other benthic species, damselfish nests or gardens, and predation by the corallivorous snail *Coralliophila abbreviata.*

Mortality as a result of predation on *D. cylindrus* by the corallivorous snail *C. abbreviata* was recorded in 26 of the 3065 observations (< 1%). The amount of recent mortality attributed to these snails on affected colonies averaged 1.1%. Mortality caused by snail predation was seen across all seasons, but spiked in November – January (Figure 7).

Recent mortality as a result of competition with boring sponges in the genus *Cliona* or competition with the encrusting octocoral *Erythropodium caribaeorum* was seen in 17 of the 3065 observations (< 1%). Average recent mortality on affected colonies was 0.7% and was observed across all seasons.

Of the 3065 observations, 16 (< 1%) identified recent mortality associated with tissue that had been abraded or buried. Two instances of tissue mortality were attributed to abrasion by marine debris (polyethylene trap rope). Average recent mortality on colonies experiencing abrasion or burial was 1.4%. These stressors were observed across all seasons (Figure 7).

### Bleaching Events

Of the 3065 total observations, 79 (2.6%) documented recent mortality attributable to bleaching; that recent mortality averaged 7.2%. In 27 observations, bleaching-related recent mortality exceeded 5%, and in 3 of those it exceeded 60%. The majority (70%) of bleaching-related mortality observations resulted from the 2014 summer hyperthermal event and were seen at eight sites from Key Largo to Key West. The remaining observations (30%) followed the 2015 hyperthermal event and were documented at two sites, one in the Upper Keys and one in the Lower Keys. More detailed information on bleaching-related patterns of Florida *D. cylindrus* colonies during these events can be found in Lewis et al. (2019).

Hourly in situ water temperatures from *D. cylindrus* sites across the Florida Keys recorded elevated summer temperatures in 2014 and 2015. The NOAA Degree Heating Week index also identified 2014 and 2015 as high thermal stress years. *D. cylindrus* bleaching events followed these warm-water periods (Figure 8). During other years (2013, 2016-2019), some paling was recorded on some colonies during the late summer months, but bleaching and/or bleaching-related mortality was not observed on *D. cylindrus*. Thermal stress, as measured by Degree Heating Weeks, was high in 2019 as well, but no bleaching beyond some instances of paling was observed on the species.

**Figure 8.**
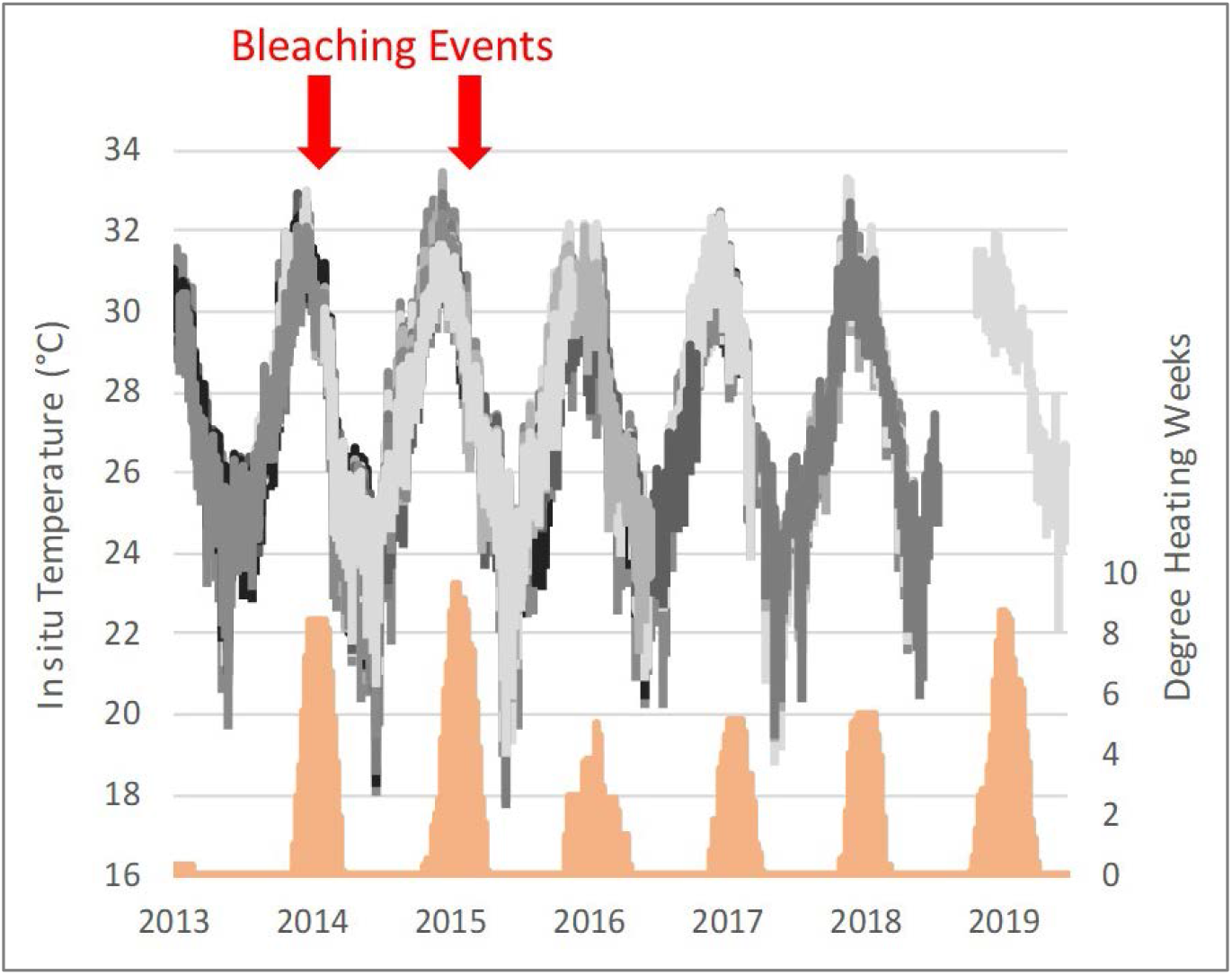
Degree heating weeks from NOAA’s Florida Keys Coral Reef Watch (orange) and hourly *in situ* temperature records from *Dendrogyra cylindrus* sites from 2012-2019. Bleaching events correspond with excessive thermal stress events in 2014 and 2015.

### Disease (excluding SCTLD)

A variety of diseases were observed on *D. cylindrus* colonies: black band disease, an unidentified yellow band disease, white plague, and stony coral tissue loss disease (SCTLD). All had deleterious effects on *D. cylindrus*.

Black band disease was identified in 28 of 3065 (< 1%) monitoring events. All of these events were observed in September through December. Average recent mortality on black-band diseased colonies was 5%. More detailed information is published in Lewis et al. (2017).

An unknown disease was observed on Upper Keys *D. cylindrus* during periods of cooler water in 2014. The disease presented similarly to yellow-band disease, with a pale/yellow band of tissue often accompanied by slow tissue loss (Figure 9). This was observed in 18 of the 3065 (< 1%) observations. The disease was documented at 8 sites, all in the Upper Keys. Observations were confined to winter/spring months, with 10 of the observations in January/February and the other 8 in late April/early May. Average recent mortality on affected colonies was less than 1%.

**Figure 9.**
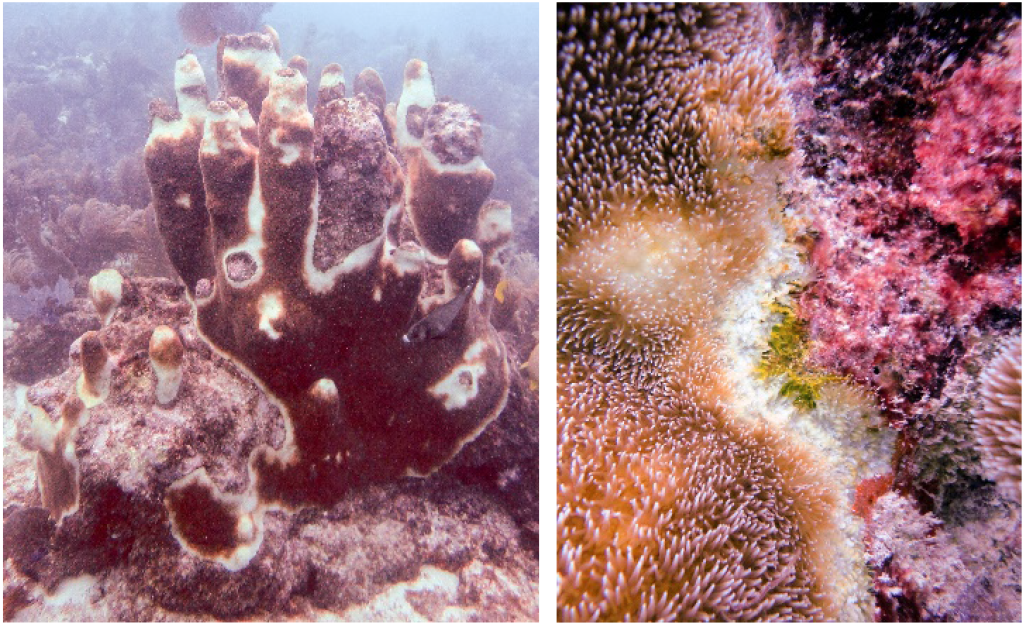
Appearance of unknown yellow band-like disease on *Dendrogyra cylindrus*. Both images are from the Upper Keys in December 2012.

White plague was the most common and deleterious disease impacting Florida *D. cylindrus* before the arrival of SCTLD (discussed below). White plague was documented in 371 of the 3056 observations (12%). Of affected colonies, the average recent mortality was 6.3%. Observations of white plague were distributed across the Florida Reef Tract. Prevalence of white plague in 2013-2015 (before the appearance of SCTLD) increased during and following periods of heat stress. Low levels of heat stress in the fall of 2013 were marked by peak white plague prevalence under 20%. High levels of heat stress in 2014 and 2015 were marked by peak white plague prevalence levels over 40% (Figure 10).

**Figure 10.**
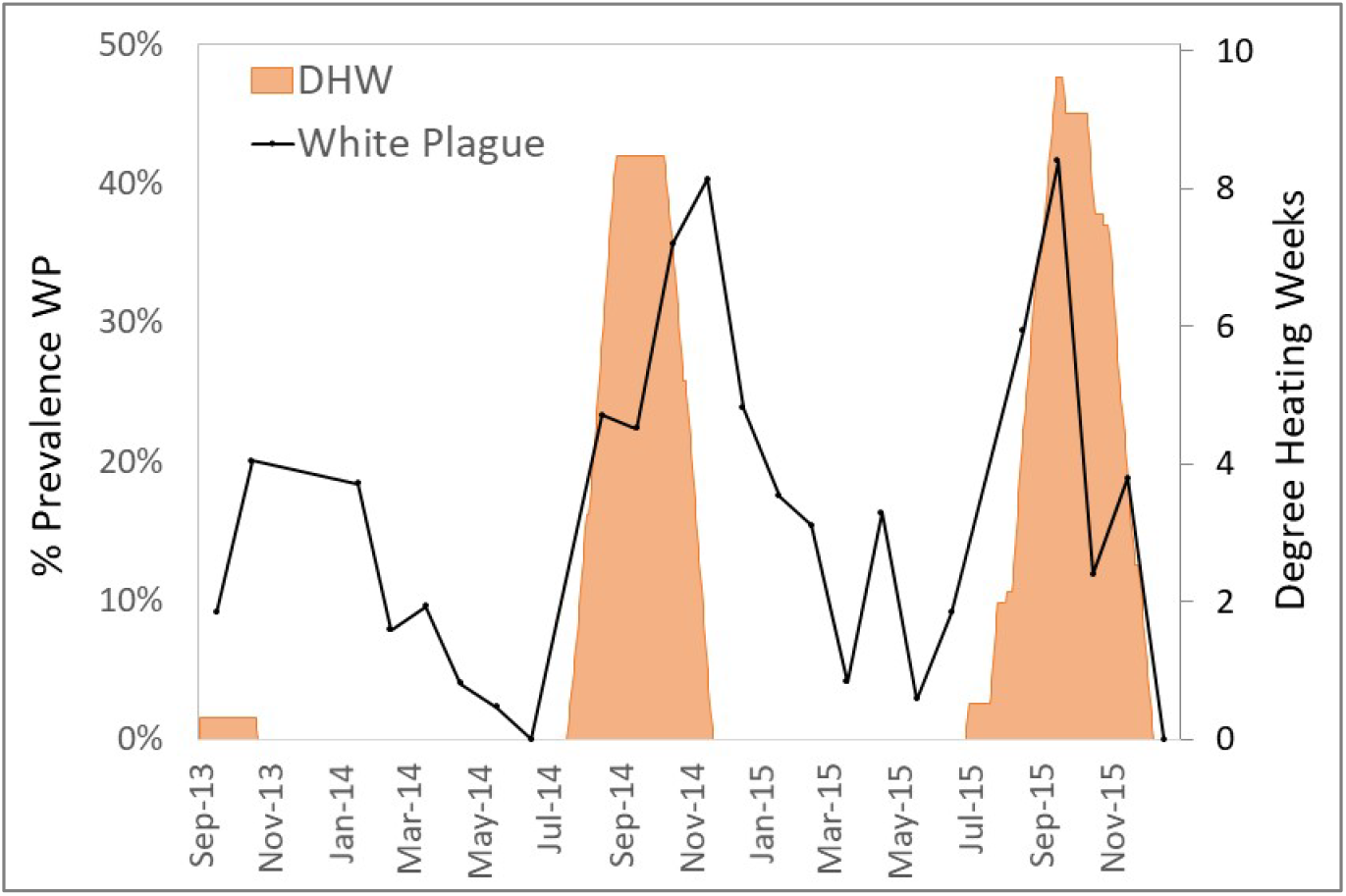
Percentage of surveyed *Dendrogyra cylindrus* with recent mortality caused by white plague during each month (solid line). Cumulative degree heating weeks (DHW), a measure of thermal stress as determined by NOAA are shown in orange fill.

### Stony Coral Tissue Loss Disease

SCTLD was identified as distinct from white plague following raised awareness from Precht et al. (2016) and subsequent regional collaborative efforts that resulted in a case definition (FKNMS/DEP 2018). Photos of colonies from the Florida Keys and Biscayne regions in 2015-2017 were reassessed to correctly separate observations of white plague and SCTLD that may have been combined during the field assessments. Disease data from southeast Florida colonies in 2015 are derived from Kabay (2016) and could not be categorized post-hoc; though SCTLD was highly likely in southeast Florida *D. cylindrus colonies*, we cannot confirm or quantify that and those data are omitted. The first SCTLD observation on Florida *D. cylindrus* occurred in February 2016 in Biscayne National Park. Infections throughout the Upper Keys population occurred throughout 2016. In the Middle Keys, SCTLD was first observed at Conch Reef in February 2017, progressed southwest through Long Key in December 2017, and reached the Sombrero Reef area in April 2018. The uppermost sites within the Lower Keys were also affected in April 2018, with southwestern progression continuing through all but the Dry Tortugas colonies by late 2019 (Figure 11).

**Figure 11.**
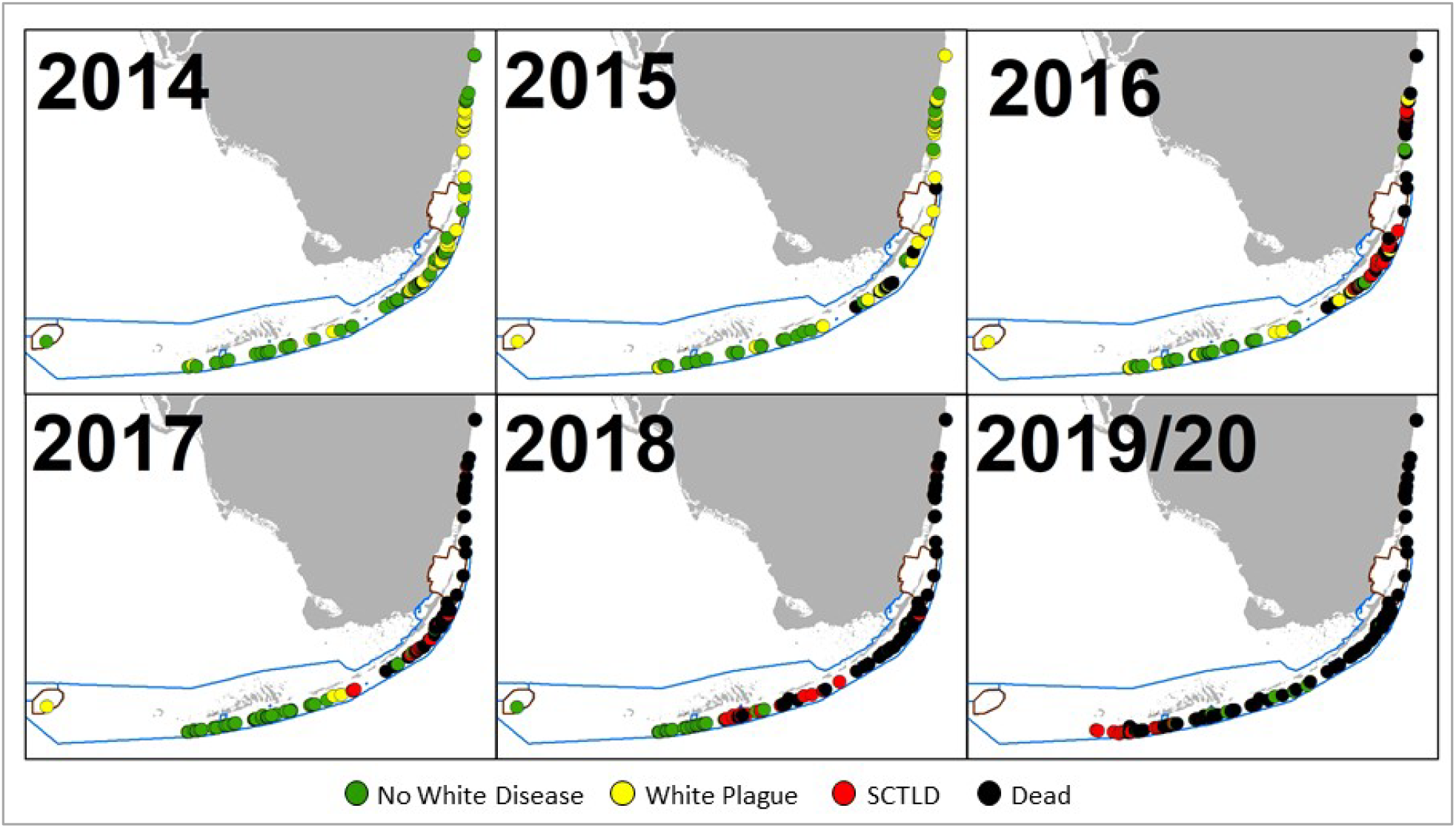
Status of *Dendrogyra cylindrus* colonies from 2014 through March 2020. At sites with multiple observations within a calendar year, the most recent is indicated. For sites with multiple colonies, disease on any colony (white plague or SCTLD) resulted in the site being classified as diseased. Stony coral tissue loss disease radiated from a site just north of Biscayne National Park and in most cases resulted in complete mortality. SCTLD was not identified until 2016, and so some observations of white plague in 2015 in southeast Florida and the Upper Keys may be SCTLD.

Quantifying the proportion of *D. cylindrus* colonies affected by SCTLD is complicated by spatial and temporal variation as the disease front progressed. Its rapid spread across infected colonies occasionally resulted in complete mortality between monitoring events leading to inconclusive causation. However, there were no colonies within the disease endemic and epidemic zones that did not experience catastrophic loss as SCTLD progressed along the FRT. In regions in which SCTLD had been present for at least one year (first observed before January 2019), 100% of colonies either a) had active SCTLD, b) were completely dead and assumed to be SCTLD-affected due to the rapid nature of the mortality, or c) suffered severe tissue loss leaving only small patches of remaining tissue. Colonies observed with active SCTLD had an average recent mortality of 19% (± 27% SD).

### Summary of Losses

Stressors varied in their prevalence (frequency of observation) and severity (% of recent mortality associated with each observation). SCTLD impacted the greatest number of *D. cylindrus* colonies, followed by white plague and bleaching-related mortality (Table 2). SCTLD also had the highest severity, followed by white plague, bleaching, and black band disease.

**Table 2.**
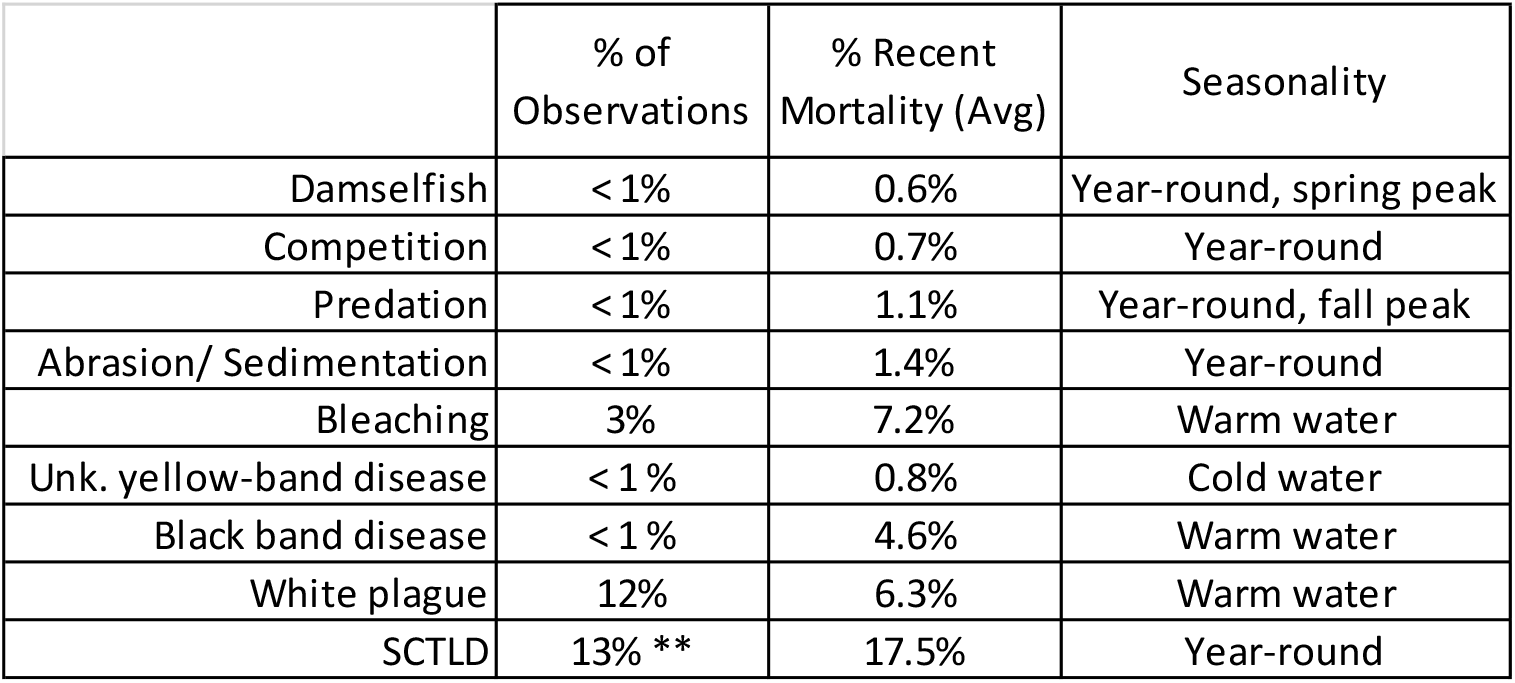
Summary of stressors affecting *D. cylindrus* colonies across 3056 observations between 2013-2019. While stressors are observable only during snapshot monitoring events, the percent of colonies affected and the percent of recent mortality due to SCTLD (**) presented here are particularly conservative as rapid mortality resulted in many colonies dying between monitoring events.

Colonies with at least one annual monitoring event for each year from 2013/14 – 2019/20 were analyzed for tissue loss over time. Between 41 and 65 colonies representing 8 to 20 genotypes from each region were assessed (Figure 12). Heavy tissue losses (24-43%) occurred across all regions between the 2013/2014 baseline and the end of 2015. Those losses were dominated by white plague and bleaching. As SCTLD reached each region, losses were rapid, generally leading to complete mortality. Within the Upper Keys/Biscayne region, the full subset of colonies was extinct by the end of 2018. Within the Middle Keys region, the amount of live tissue stabilized during 2016, but SCTLD impacts from 2017 through 2019 resulted in a 97.4% loss of tissue by March 2020. In the Lower Keys the amount of tissue was stable through 2017 and 2018 (despite the direct overpass of a category 4 hurricane), but the arrival of SCTLD in 2018 led to a 70% loss of tissue through March 2020 (Figure 12).

**Figure 12.**
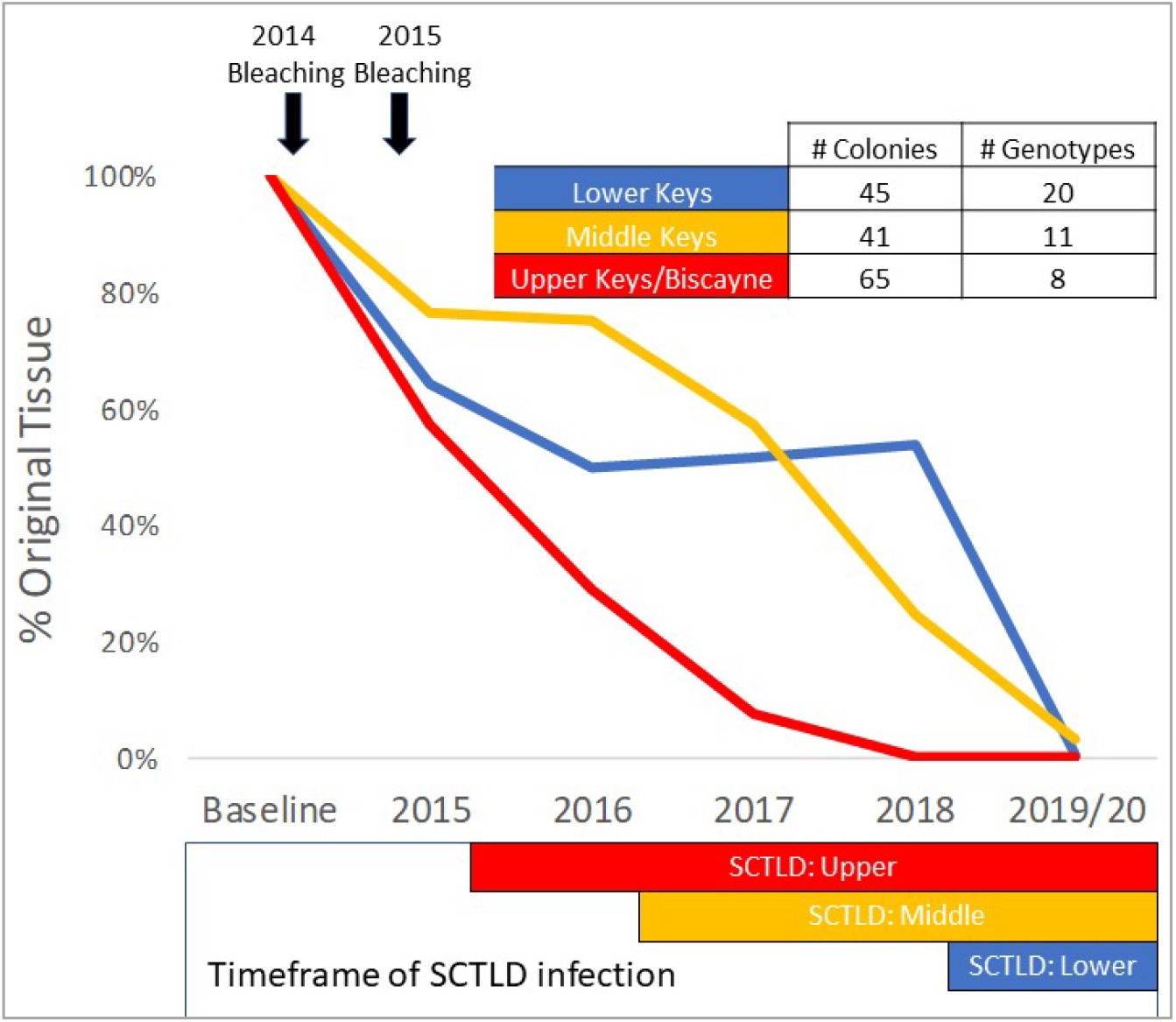
Percent of original tissue on a subset of *Dendrogyra cylindrus* colonies with at least one annual monitoring from 2013/14 through March 2020. Total tissue surface area (cm^2^) is calculated from size measurements and percent live coral. Amount of tissue each year on the selected colonies within each region was summed and compared to baseline surveys. The timing of the 2014 and 2015 bleaching events is highlighted (top), as is the presence of Stony Coral Tissue Loss Disease within each region (bottom).

### Population status in March 2020

Extinction losses of *D. cylindrus* on the Florida Reef Tract from the baseline 2013/14 surveys through March 2020 are as follows:

- 73% loss of genotypes (not including the 13% with active SCTLD in March 2020)
- 88% loss of colonies
- 91% loss of tissue

In March 2020, 50 genotypes remained. Of these, 25 were actively experiencing rapid tissue loss due to stony coral tissue loss disease, 5 had been treated with topical antibiotics to arrest SCTLD, and 10 had declined to less than 2% coral cover since the baseline surveys. Of the remaining 10 healthy and intact genotypes, 9 were located on the far western edge of the Florida Reef Tract where SCTLD had not yet reached.

## Discussion

### Demographics & Genetic Diversity

Demographic surveys of Florida Reef Tract *D. cylindrus* colonies indicate a population comprised of predominantly large and presumably old individuals. Most small colonies were very near larger colonies and had the appearance of fragmented pillars. This project documented only one small colony (< 20 cm) that was greater than 70m from another colony and presumed to be a sexual recruit. Similarly, ongoing Florida monitoring programs, which survey hundreds of sites per year, have not documented juvenile *D. cylindrus* (CREMP pers comm; Miller (2000-2011)). Growth rates of *D. cylindrus* are poorly understood, but measurements of transplanted pillars in Key Largo from 1993-1995 recorded an average vertical pillar extension of 1.8 ± 0.3 SD cm per year (Hudson and Goodwin 1997). Making the liberal assumptions that the same growth rate exists in the development of the base in vertical and horizontal directions and that the rate is constant from settlement onwards, only 11 of the FRT genotypes (6%) could potentially be younger than 20 years. Of note, one small colony (46 cm x 43 cm) without any vertical pillar development was found on an artificial substrate sunk in 1986 (FWC/DMF 2020), indicating a maximum age of 31 years when first measured.

Demographic surveys conducted on a *D. cylindrus* population in Colombia (Acosta and Acevedo 2006) allow for comparisons between regions. The Florida population has significantly larger colonies (Mann-Whitney Rank Sum Test p < 0.001; Figure 13). In the Colombian study, the population was comprised mostly (52%) of colonies less than 60 cm in height. In contrast, less than 20% of Florida colonies fell within that size class. In other coral species, such a lack of juveniles has been suggested to correlate with poor recruitment resulting from reef degradation (Meesters et al. 2001).

**Figure 13.**
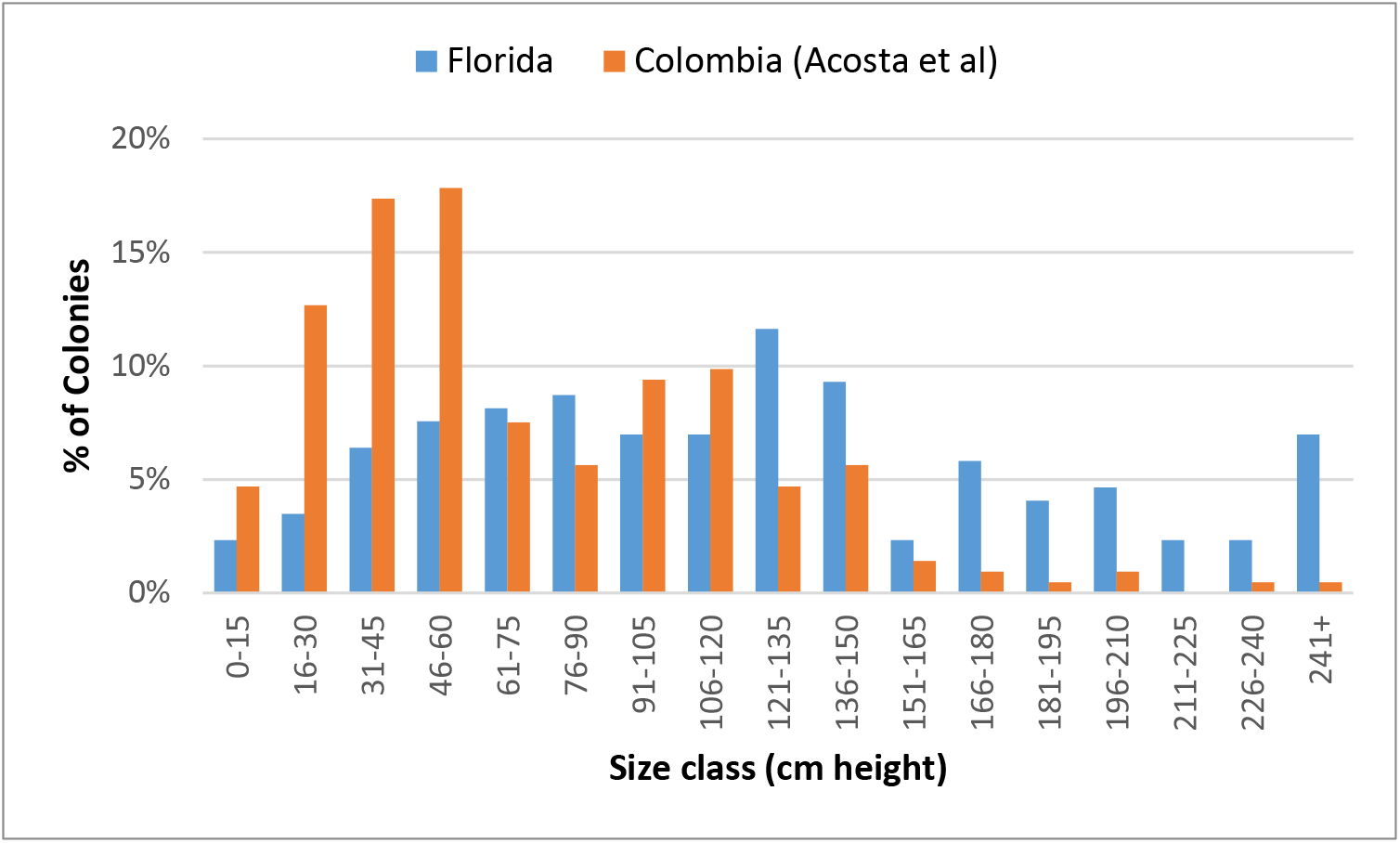
Demographic comparisons of *Dendrogyra cylindrus* height distributions in Florida and Colombia. Colombian data taken from Acosta and Acevedo, 2006.

The near absence of juvenile *D. cylindrus* in Florida may be a result of a) poor juvenile survival, b) poor settlement rates, and/or c) poor fertilization. We will examine each of these in turn based on the population demographics.

If low juvenile survival were causing the lack of observed juveniles, mortality would need to occur quite early to escape notice by various monitoring programs. The fixed-site and stratified random monitoring programs that have noted an absence of juveniles operate by recording all corals greater than 4 cm in diameter, and so mortality would have to occur before that size. While growth rates of juvenile *D. cylindrus* settlers have never been quantified in the wild, larvae settled in laboratory settings have remained smaller than 2 cm for up to three years, which in the wild could be overlooked. However, recruitment studies conducted throughout the Florida Keys which focus on new settlers have also never documented juvenile *D. cylindrus* (CREMP, pers comm; Bartlett et al. (2018)).

Poor larval settlement may be a possible bottleneck leading to the absence of juveniles. Coral recruitment rates in Florida were identified by Moulding (2005) to be highest in the Lower Keys, with decreasing recruitment in the Middle and Upper Keys. The smaller average size of *D. cylindrus* colonies in the Lower Keys could be a result of more common or more recently viable recruitment within that region. However, very little is known about settlement cues, growth rates, or mortality curves for *D. cylindrus*, and this hypothesis would require further testing.

The third hypothesis is that the number of larvae available for settlement is limited. Corals are generally considered r-selected species; millions of eggs are produced, and survival is extremely low through the early life stages. In the case of Florida’s *D. cylindrus* population, fertilization is likely to be essentially non-existent due to several factors:

1. Of the 188 genotypes, only 4 were within 10 meters of a genetically different neighbor. Diffusion and fertilization across even 10 meters is unlikely given the obstacles of predation, physical barriers, and currents.
2. In contrast to most broadcast spawners that have positively buoyant gametes, eggs and sperm of *D. cylindrus* do not rapidly rise to the surface. Lab and field observations indicate that eggs are only slightly positively buoyant, while sperm is neutral or slightly negative. As such, fertilization success requires gametes navigating three dimensions rather than mixing in surface slicks, which adds an additional barrier to fertilization potential.
3. *Dendrogyra cylindrus* colonies are mostly gonochoric (but see Neely et al. (2018)). Thus, even if gametes from genetically-distinct nearby colonies do find each other, there is only a 50% chance of gender compatibility if 1:1 sex ratios are assumed.

Other Caribbean populations of *D. cylindrus* in which multiple genotypes are in close proximity to each other are likely to overcome these obstacles. However, the population on the Florida Reef Tract, even before the losses of 2014-2020, was largely constrained by Allee effects that appear to have rendered the population reproductively extinct for decades.

### Sources of Mortality

*Dendrogyra cylindrus* colonies in Florida were subjected to a variety of stressors between 2014 and early 2020 that resulted in tissue loss and colony mortality. These included chronic and relatively low-impact factors such as predation by *Coralliophila abbreviata*, damselfish gardens and nests, competition with other benthic organisms, and abrasion or tissue burial. These stressors were uncommon (< 1 % of observations) and minor (≤ 1% recent mortality). Though not quantified, we observed many of these minor sources of mortality healing with tissue regrowth in subsequent surveys. We suggest that losses from these minor, chronic stressors can be balanced by tissue regrowth and do not necessarily lead to net tissue loss over time.

Acute stressors were more prevalent and detrimental. Four different diseases impacted *D. cylindrus* populations during the survey period: an unidentified yellow-band disease, black band disease, white plague, and SCTLD. These diseases varied in prevalence, seasonality, and resulting mortality. Black-band disease and white plague are both known in other species and regions to increase in prevalence during seasonal hyperthermal events (Kuta and Richardson 1996; Cróquer and Weil 2009). Before the emergence of SCTLD, white plague had the greatest impact on the Florida population, particularly during warm-water events, and contributed to unsustainable losses. White plague was documented during 12% of observations, with seasonal peaks of 40% and 42% prevalence in 2014 and 2015 respectively. In comparison, observations on a *D. cylindrus* population in Colombia documented white disease incidence of 9.4% in January 2002 and 1.5% in October 2012 (Bernal-Sotelo et al. 2019).

The Florida *D. cylindrus* population was also heavily impacted by hyperthermal bleaching events in 2014 and 2015 that resulted in high tissue mortality at some sites. In the 1983 Florida mass thermal bleaching, Jaap (1985) identified *D. cylindrus* as bleaching resistant. Maximum water temperatures in the Lower Keys region of study during that event were 32.3°C. During the 2014 and 2015 bleaching events, in situ water temperatures in the same region exceeded 32.3° C for three and four days respectively. Single-day high temperatures are poor predictors of bleaching compared to compounded thermal stress as identified by Degree Heating Weeks (Kayanne 2017), but high thermal stress or thermal stress compounded by other stressors such as ultraviolet radiation (Gleason and Wellington 1993) or degraded water quality (Lapointe et al. 2019) in 2014 – 2015 resulted in severe bleaching and associated mortality in a species historically thought to be bleaching-resistant.

Stressors documented from the baseline surveys through the end of 2015, predominantly white plague and two bleaching events, led to substantial losses of *D. cylindrus,* including the extinction of two genotypes. However, in the Middle Keys and Lower Keys, tissue losses subsequently stabilized until SCTLD began impacting the regions. From the end of 2016 to the end of 2018, the amount of tissue on the colonies in the Lower Keys increased ~4% per year. Thus, in the absence of severe stressors, regeneration of lost tissue on extant colonies began rebuilding the population. Based on regeneration rates from the Lower Keys region, an 11 year interval would be required to replace the tissue losses that occurred as a result of the 2014-2015 mortality events. These values are similar to those predicted for the branching coral *Acropora palmata* in the Florida Keys, which suggested an 11 year recovery period following mortality from the 2005 hurricane season (Williams and Miller 2012). Many factors are likely to influence the recovery rates of colonies, including the severity and pattern of tissue mortality, the capacity of the denuded skeleton to support recolonization, and the reef conditions suitable for coral growth (Lester and Bak 1985). However, if colonies retain some live tissue, the Lower Keys data suggest that colony recovery is possible if suitable conditions extend for long enough between acute mortality events.

Though recovery may be possible in suitable conditions, the intervals between acute mortality events, including hurricanes, bleaching events, and disease outbreaks, are currently shorter than the expected recovery time. Hurricane strikes are irregular, but based on historical data, the interval of a strike to the Key West region averages every eight years (Blake et al. 2011). Lower Keys *D. cylindrus* colonies impacted by Hurricane Irma in 2017 exhibited some pillar breakage as well as lateral colony movements of up to 12 meters. However, tissue loss resulting from these processes was not substantial. The extensive clonality of *D. cylindrus* colonies within sites, including areas with up to 150 clonal ramets, suggests that this species relies on asexual fragmentation as a form of colony production and that storm-related breakage can be as much a positive as a negative for the species’ success.

Other stressors that resulted in substantial tissue loss to the Florida *D. cylindrus* population are predicted to increase in frequency. Mass coral bleaching, which led to tissue and colony losses in 2014 and 2015, is expected to become an annual event across much of the Florida Reef Tract by 2040 (van Hooidonk et al. 2016). Though bleaching does not necessarily result in mortality, and some *D. cylindrus* have shown adaptability to bleaching through shuffled symbionts (Lewis et al. 2019), recovery of typical symbiont and microbial communities may take several years (Edmunds 2014, Kemp et al 2014, Bourne 2007). Due to sub-optimal holobiont partnerships, bleaching stress is likely to alter long-term physiological functions (Ban et al. 2014), including reduced colony growth and reproductive output (Mendes and Woodley 2002).

Disease has been the primary driver of population decline in Florida *D. cylindrus.* Stony coral tissue loss disease has been unprecedented in its temporal and spatial extent and is unusual in maintaining its virulence throughout multiple seasonal cycles. Prior to SCTLD, *D. cylindrus* colonies were mostly affected by white plague and black band disease, both associated with high thermal stress. This mirrors correlations between hyperthermal stress and disease prevalence in other regions and species (Cróquer and Weil 2009). Modelling by Maynard et al. (2015) projected the timing of disease conditions using host susceptibility, pathogen abundance, and pathogen virulence. In Florida, these models show that at least two of the three disease-enhancing scenarios will be met approximately 25 years before annual bleaching is predicted. With van Hooidonk et al. (2016) predicting annual bleaching throughout southeast Florida in 2040, this 25-year earlier disease prediction coincided with the rise of SCTLD and widespread loss of *D. cylindrus* to disease.

Model predictions of future reef stressors are largely irrelevant for the Florida *D. cylindrus* population. Initial small population size and geographic distances between genotypes most likely rendered the Florida population reproductively extinct during the 20^th^ century. Genetic analyses by Chan et al. (2019) indicate that the Florida *D. cylindrus* population is distinct from other Caribbean populations, so influx from other regions is neither historically significant nor likely to be an option for natural replenishment. The Florida colonies have suffered catastrophic loss, with the most recent and devastating cause of mortality being SCTLD. Losses in genetic diversity, colony numbers, and live tissue have rendered the species functionally extinct on the FRT, with no ability for natural recovery.

### Future of the Species

In Florida, *D. cylindrus* is reproductively and functionally extinct. The few remaining bits of live tissue are small, geographically isolated, and vulnerable to chronic stressors as well as widespread acute stressors such as SCTLD infection or the next thermally-associated bleaching and disease events.

In response to *D. cylindrus* losses in 2014 and 2015, a genetic rescue program was initiated in late 2015. Fragments were removed from the wild for safekeeping at nursery and land-based facilities. Through this effort, the extinction of Florida genotypes was halted by maintaining genetic material and creating grow-out and brood stock. At the end of 2019, 543 fragments representing 123 genotypes were held in land-based and offshore nurseries. In addition to maintaining the Florida genetic stock, these fragments have also been used as the broodstock in the first successful controlled reproduction of an Atlantic species (Johnson 2019) which has increased the genetic diversity of the population within the genetic bank. Much like the California condor (Wallace and Toone 1992), the fate of *D. cylindrus* lies in captive breeding to produce diverse as well as resilient genotypes, Large-scale efforts to improve water quality and curb climate change are also essential for creating the conditions that will allow for the successful future restoration, survival, and wild reproduction of this iconic and unique coral.

## Acknowledgements

We thank scientific divers from Florida Aquarium, Keys Marine Laboratory, Florida FWC/FWRI, Florida International University, and Nova Southeastern University for assistance with fieldwork. We are additionally grateful to individuals, dive shops, and research programs that provided site locations. This research was supported by Florida’s Wildlife Legacy Initiative (State Wildlife Grants CFDA No. 15.634), the United States Fish and Wildlife Service grants program, Marine Projects Grant Cycle (federal award No. F13AF01085), National Science Foundation Rapid Response Research grant #1503483, the Florida Aquarium’s Center for Conservation, and the Florida International University Integrative Marine Genomics and Symbiosis (IMaGeS) lab. The work was conducted under Florida Keys National Marine Sanctuary permits FKNMS-2013-085-A1, FKNMS-2014-004-A1, FKNMS-2016-004-A1, Dry Tortugas Permit DRTO-2015-SCI-0018, Biscayne National Park Permits BISC-2013-SCI-018 and BISC-2015-SCI-0019, and State of Florida permit SAL-13-1451-SRP.

## Literature Cited

Acosta A, Acevedo A (2006) Population structure and colony condition of *Dendrogyra cylindrus* (Anthozoa: Scleractinea) in Providencia Island, Colombian Caribbean. Proceedings of the 10th International Coral Reef Symposium 10:1605–1610

Aronson R, Bruckner A, Moore JA, Precht B, Weil E (2008) Dendrogyra cylindrus. The IUCN Red List of Threatened Species e T133124A3582471, https://dx.doi.org/10.2305/IUCN.UK.2008.RLTS.T133124A3582471.en.

Aronson RB, Precht WF (2001) White-band disease and the changing face of Caribbean coral reefs. Hydrobiologia 460:25–38

Bartlett LA, Brinkhuis VI, Ruzicka RR, Colella MA, Lunz KS, Leone EH, Hallock P (2018) Dynamics of stony coral and octocoral juvenile assemblages following disturbance on patch reefs of the Florida Reef Tract.) Corals in a changing world IntechOpen, London:99–120

Baums IB, Miller MW, Szmant AM (2003) Ecology of a corallivorous gastropod, *Coralliophila abbreviata*, on two scleractinian hosts. 1: Population structure of snails and corals. Mar Biol 142:1083–1091

Bernal-Sotelo K, Acosta A, Cortés J (2019) Decadal Change in the Population of *Dendrogyra cylindrus* (Scleractinia: Meandrinidae) in Old Providence and St. Catalina Islands, Colombian Caribbean. Frontiers in Marine Science 5

Blake ES, Landsea CW, Gibney EJ (2011) The deadliest, costliest, and most intense United States tropical cyclones from 1851 to 2010 (and other frequently requested hurricane facts) NOAA Technical Memorandum NSW NHC-6. NOAA/NWS/NCEP/National Hurricane Center

Causey B (2008) Coral Reefs of the U.S. Caribbean: The History of Massive Coral Bleaching and other Perturbations in the Florida Keys. In: Wilkinson C, Souter D (eds) Status of Caribbean coral reefs after bleaching and hurricanes in 2005. Global Coral Reef Monitoring Network, and Reef and Rainforest Research Center, Townsville, QLD, pp 61–67

Chan AN, Lewis CL, Neely KL, Baums IB (2019) Fallen Pillars: The Past, Present, and Future Population Dynamics of a Rare, Specialist Coral–Algal Symbiosis. Frontiers in Marine Science 6

Courchamp F, Clutton-Brock T, Grenfell B (1999) Inverse density dependence and the Allee effect. Trends Ecol Evol 14:405–410

Cróquer A, Weil E (2009) Changes in Caribbean coral disease prevalence after the 2005 bleaching event. Diseases of aquatic organisms 87:33–43

FKNMS/DEP (2018) Case Definition: Stony Coral Tissue Loss Disease, https://floridadep.gov/sites/default/files/Copy%20of%20StonyCoralTissueLossDiseaseCaseDefinition%20final%2010022018.pdf

FRRP (2015) Florida Reef Resilience Program’s Disturbance Response Monitoring Florida Keys Reef Zones GIS Shapefile (Version 2)

FWC/DMF (2020) Artificial Reefs Florida. FWC, DMF, http://myfwc.com/conservation/saltwater/artificial-reefs/locate-reefs/

FWRI (2014) Unified Florida Coral Reef Tract Map v1.2

Gleason DF, Wellington GM (1993) Ultraviolet Radiation and Coral Bleaching. Nature 365:836–838

Goreau TJ, Cervino JM, Goreau M, Hayes R, Hayes M, Richardson L, Smith G, DeMeyer K, Nagelkerken I, Garzon-Ferreira J, Gil D, Garrison G, Williams EH, Bunkley-Williams L, Quirolo C, Patterson K, Porter JW, Porter K (1998) Rapid spread of diseases in Caribbean coral reefs. Rev Biol Trop 46:157–171

Hudson JH, Goodwin WB (1997) Restoration and growth rate of hurricane damaged pillar coral *(Dendrogyra cylindrus)* in the Key Largo National Marine Sanctuary, Florida. Proceedings of the 8th International Coral Reef Symposium 1:567–570

Jaap W (1985) An epidemic zooxanthellae expulsion during 1983 in the lower Florida Keys coral reefs: hyperthermic etiology. Proc 5th Int Coral Reef Symp 6:143–148

Johnson LM (2019) A scientific breakthrough at the Florida Aquarium could save “America’s Great Barrier Reef”, CNN

Kabay L (2016) Population Demographics and Sexual Reproduction Potential of the Pillar Coral, Dendrogyra cylindrus, on the Florida Reef Tract. Nova Southeastern University,

Kayanne H (2017) Validation of degree heating weeks as a coral bleaching index in the northwestern Pacific. Coral Reefs 36:63–70

Kemp D, Fitt W, Schmidt G (2008) A microsampling method for genotyping coral symbionts. Coral Reefs 27:289–293

Kuta K, Richardson L (1996) Abundance and distribution of black band disease on coral reefs in the northern Florida Keys. Coral Reefs 15:219–223

Lapointe BE, Brewton RA, Herren LW, Porter JW, Hu C (2019) Nitrogen enrichment, altered stoichiometry, and coral reef decline at Looe Key, Florida Keys, USA: a 3-decade study. Mar Biol 166:108

Lester RT, Bak RPM (1985) Effects of environment on regeneration rate of tissue lesions in the reef coral *Montastrea annularis* (Scleractinia). Mar Ecol Prog Ser 24:183–185

Lewis C, Neely K, Rodriguez-Lanetty M (2019) Recurring Episodes of Thermal Stress Shift the Balance From a Dominant Host-Specialist to a Background Host-Generalist Zooxanthella in the Threatened Pillar Coral, *Dendrogyra cylindrus*. Frontiers in Marine Science 6

Lewis CL, Neely KL, Richardson LL, Rodriguez-Lanetty M (2017) Temporal dynamics of black band disease affecting pillar coral *(Dendrogyra cylindrus)* following two consecutive hyperthermal events on the Florida Reef Tract. Coral Reefs 36:427–431

Maynard J, van Hooidonk R, Eakin CM, Puotinen M, Garren M, Williams G, Heron SF, Lamb J, Weil E, Willis B, Harvell CD (2015) Projections of climate conditions that increase coral disease susceptibility and pathogen abundance and virulence. Nature Climate Change 5:688–694

Meesters EH, Hilterman M, Kardinaal E, Keetman M, de Vries M, Bak RPM (2001) Colony size-frequency distributions of scleractinian coral populations: spatial and interspecific variation. Mar Ecol Prog Ser 209:43–54

Mendes JM, Woodley JD (2002) Effect of the 1995-1996 bleaching event on polyp tissue depth, growth, reproduction and skeletal band formation in *Montastraea annularis*. Mar Ecol Prog Ser 235:93–102

Miller SM (2000-2011) Quick Look Reports and Data Summaries, CMS/UINCW, Key Largo FL

Moulding A (2005) Coral recruitment patterns in the Florida Keys. Rev Biol Trop 53 Suppl 1:75–82

Neely KL, Lewis C, Chan A, Baums IB (2018) Hermaphroditic spawning by the gonochoric pillar coral *Dendrogyra cylindrus*. Coral Reefs 37:1087–1092

NOAA (2013-2019) NOAA Coral Reef Watch Version 3.1 Daily GLobal 5-km Satellite Coral Bleaching Degree Heating Week Product, College Park, MD, USA

Porter JW, Meier OW (1992) Quantification of Loss and Change in Floridian Reef Coral Populations. Am Zool 32:625–640

Precht WF, Gintert BE, Robbart ML, Fura R, van Woesik R (2016) Unprecedented Disease-Related Coral Mortality in Southeastern Florida. Scientific Reports 6

Richardson L, Goldberg W, Carlton RG, Halas JC (1998) Coral disease outbreak in the Florida Keys: Plague Type II

Ruzicka RR, Colella MA, Porter JW, Morrison JM, Kidney JA, Brinkhuis V, Lunz KS, Macaulay KA, Bartlett LA, Meyers MK, Colee J (2013) Temporal changes in benthic assemblages on Florida Keys reefs 11 years after the 1997/1998 El Nino. Mar Ecol Prog Ser 489:125–141

Sutherland KP, Porter JW, Torres C (2004) Disease and immunity in Caribbean and Indo-Pacific zooxanthellate corals. Mar Ecol Prog Ser 266:273–302

Szmant AM (1986) Reproductive Ecology of Caribbean Reef Corals. Coral Reefs 5:43–53

Thurber RLV, Burkepile DE, Fuchs C, Shantz AA, McMinds R, Zaneveld JR (2014) Chronic nutrient enrichment increases prevalence and severity of coral disease and bleaching. Glob Change Biol 20:544–554

USFWS (2014) Endangered and Threatened Wildlife and Plants: Adding 20 Coral Species ot the List of Endangerd and Threatened Wildlife: 50 CFR Part 17. Federal Registry 59:67356–67359

van Hooidonk R, Maynard J, Tamelander J, Gove J, Ahmadia G, Raymundo L, Williams G, Heron SF, Planes S (2016) Local-scale projections of coral reef futures and implications of the Paris Agreement. Scientific Reports 6

Wallace M, Toone W (1992) Captive management for the long term survival of the California condor Wildlife 2001: populations. Springer, pp 766–774

Williams DE, Miller MW (2012) Attributing mortality among drivers of population decline in *Acropora palmata* in the Florida Keys (USA). Coral Reefs 31:369–382

